# Real-time feedback reduces participant motion during task-based fMRI

**DOI:** 10.1101/2023.01.12.523791

**Authors:** Chad S. Rogers, Michael S. Jones, Sarah McConkey, Drew J. McLaughlin, Jonathan E. Peelle

## Abstract

The potential negative impact of head movement during fMRI has long been appreciated. Although a variety of prospective and retrospective approaches have been developed to help mitigate these effects, reducing head movement in the first place remains the most appealing strategy for optimizing data quality. Real-time interventions, in which participants are provided feedback regarding their scan-to-scan motion, have recently shown promise in reducing motion during resting state fMRI. However, whether feedback might similarly reduce motion during task-based fMRI is an open question. In particular, it is unclear whether participants can effectively monitor motion feedback while attending to task-related demands. Here we assessed whether a combination of real-time and between-run feedback could reduce head motion during task-based fMRI. During an auditory word repetition task, 78 adult participants (aged 19–81) were pseudorandomly assigned to receive feedback or not. Feedback was provided FIRMM software that used real-time calculation of realignment parameters to estimate participant motion. We quantified movement using framewise displacement (FD). We found that motion feedback resulted in a statistically significant reduction in participant head motion, with a small-to-moderate effect size (reducing average FD from 0.347 to 0.282). Reductions were most apparent in high-motion events. We conclude that under some circumstances real-time feedback may reduce head motion during task-based fMRI, although its effectiveness may depend on the specific participant population and task demands of a given study.

## Introduction

Head movement during fMRI can introduce a number of different challenges to data analysis. Although long appreciated (Friston et al., 1996), the detrimental influence of head motion on resting state functional connectivity has only recently been widely recognized (Power et al., 2015). Rigid body realignment—a mainstay of fMRI analysis for decades—goes some way towards improving correspondence across images (Ashburner and Friston, 2004), but does not remove extraneous signal components introduced by movement (Friston et al., 1996). A common approach for mitigating motion-related artifacts is to include the 6 realignment parameters (translation and rotation around the X, Y, and Z axes) as nuisance regressors in first-level models. Additional approaches include wavelet despiking (Patel et al., 2014), ICA (Pruim et al., 2015), robust-weighted least squares (Diedrichsen and Shadmehr, 2005), Bayesian approaches (Eklund et al., 2017), and frame censoring (Lemieux et al., 2007). However, recent work investigating motion correction strategies in multiple data sets suggests that the optimal strategy may depend on the specific data set and output metric (Jones et al., 2022), precluding a simple one-size-fits-all solution.

Given the lack of certainty regarding how to best handle head motion during analysis, reducing head motion in the first place is a particularly appealing option. Common approaches to reducing head motion during MRI scanning include foam padding and instructing participants to not move during the scan. Other existing strategies include using custom head molds (Power et al., 2019) and playing participants a relaxing abstract movie (Vanderwal et al., 2015).

A complementary approach for reducing head motion is to provide participants with feedback regarding their head movement so they can learn to minimize motion. FIRMM software^1^ (Dosenbach et al., 2017) is one implementation of motion feedback. FIRMM uses rapid image reconstruction and rigid-body alignment to estimate frame-by-frame movement, providing visual cues to participants based on estimated movement. Feedback can either be provided in real time, or between scanning runs. FIRMM has been shown to reduce movement during resting state scans in both adults (Dosenbach et al., 2017) and in young children (Greene et al., 2018; Badke D’Andrea et al., 2022).

Despite its success in resting state fMRI, it is unclear whether motion feedback would lead to similar improvements during a typical task-based fMRI paradigm. Task-based paradigms have been shown to produce greater head motion both in terms of displacement and rotation relative to resting state scans (Huang, 2019), and could represent a natural target for motion feedback. On the one hand, if participants are able to modulate their motion in one experimental paradigm (namely, resting state studies) it stands to reason they could do so in a different experimental context (a task-based study). On the other hand, task-based fMRI studies require participants to attend to the experimental task at hand, which will reduce the cognitive resources available for noting and processing the feedback, as well as the ability to attend to proprioceptive cues related to self-motion. In addition, if experimental stimuli and motion feedback are in the same modality, subjects may also have difficulty accurately processing motion-related feedback.

Most fMRI processing pipelines rely on 6 parameter rigid body procedures to correct for frame-to-frame movement. The estimated translations in the x, y, and z planes (and rotations around the x, y, and z axes) provide 6 parameters that are typically used as a measurement of motion. Framewise displacement (FD) has been increasingly used to summarize total motion across these 6 parameters (with rotations converted to distance using simplifying assumptions about head shape) (Power et al., 2012). FD relies on absolute values of differences and thus is always positive; larger numbers reflect more total movement.

To evaluate the degree to which feedback might affect head motion during task-based fMRI, we assigned participants in a task-based fMRI study of spoken word recognition to receive feedback (the feedback group) or not (the no-feedback group). All other instructions, scanning parameters, and task requirements were identical. We included both young and older adults, and a task (word repetition) that we anticipated would typically require head movement. Finally, we were able to capitalize on the fact that each subject has many observations (hundreds of frames) by using linear mixed effects analysis.

## Method

MRI data are available from OpenNeuro (Markiewicz et al., 2021) at https://doi.org/10.18112/openneuro.ds004285.v1.0.0, reference number ds004285, and motion data and analysis scripts are available from https://github.com/jpeelle/motion-feedback. (Although we collected these data in the context of a task-based fMRI study, described below, here we focus only on effects of feedback on head motion and not the task results.)

### Participants

We tested 78 participants aged 19–81. All reported themselves to have normal hearing, to be right-handed, and not have a history of neurological disease. There were no exclusions based on sex, gender, race, or ethnicity. Participants were limited to adults because of our scientific focus on adult language processing. Because the task involved spoken language, all were native speakers of American English.

Participants were randomly assigned to control (no motion feedback) or intervention (real-time motion feedback) groups. Demographic characteristics for these groups are shown in **Table 1**. Because data collection was ended early due to COVID-19 safety requirements the groups did not have equal numbers of participants. All provided informed consent under a process approved by the Washington University in Saint Louis Institutional Review Board.

**Table 1.** Participant characteristics.

|  |  | No feedback | Feedback |
| --- | --- | --- | --- |
| Young adults | N | 17 | 31 |
| | Age (mean ± SD) | 23.3 (3.06) | 22.4 (2.64) |
|  | Sex | 13 F, 4 M | 28 F, 3 M |
| Older adults | N | 5 | 25 |
| | Age (mean ± SD) | 70.4 (5.22) | 71.3 (3.81) |
|  | Sex | 5 F, 0 M | 17 F, 8 M |

### Procedure

The procedure is modeled after that in Rogers et al. (2020) and illustrated in **Figure 1**. Participants performed a word repetition task in which on every trial they heard a spoken word, presented in stationary background noise (3 dB SNR), and repeated it back aloud.

**Figure 1.**
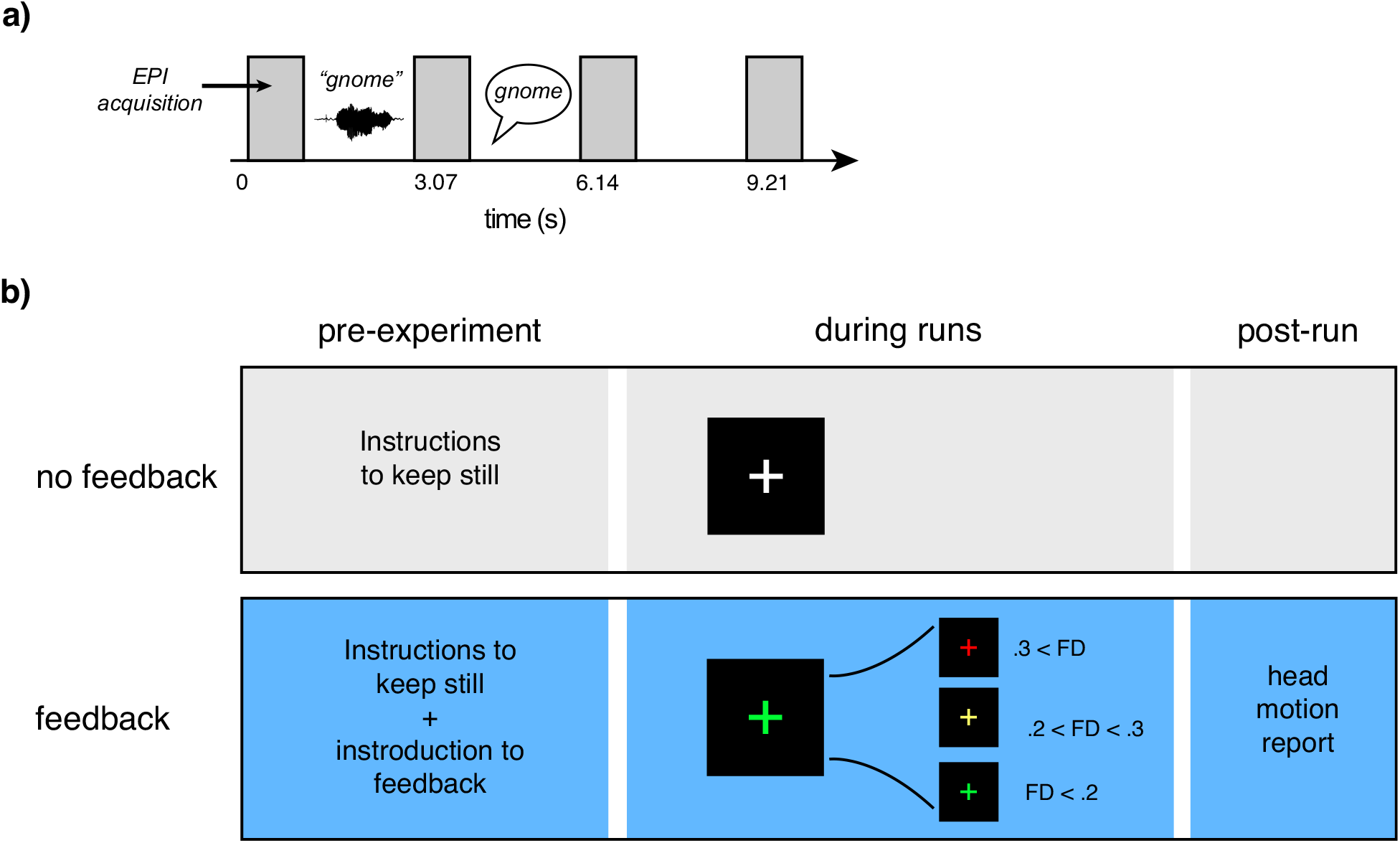
Summary of task design and processing pipeline. **a)** Schematic of the word repetition task. Participants heard a word presented in the midst of background noise during the gap between volume acquisitions; during the following gap, they repeated the word aloud. **b)** Illustration of the two groups to which a participant could be pseudorandomly assigned. The group without feedback received standard instructions to keep still at the beginning of the study; the feedback group received instructions regarding real-time feedback, a colored crosshair indicating their scan-to-scan motion (framewise displacement; FD) during each run; and a visual report on feedback at the end of each run. **c)** Illustration of time variables available for modeling.

The no-feedback group was given the following instructions:

> During this task, it is important that you hold your body and head very still. Please stay relaxed, stay alert, and keep your eyes open and on the fixation cross.

FIRMM was run in the background for experimenters to monitor motion during scanning sequences and participants were given verbal feedback after each sequence.

The group receiving motion feedback was given the following instructions:

> It is very important to remain still during your MRI so that we can obtain clear images. Even very small movements that you are not aware of can affect the image quality. While performing the task, you will be receiving feedback corresponding to your ability to remain still. This is to help you be aware of any movements you may be making. You will see a white fixation cross on the screen. The cross will change to yellow and then red depending on how much you are moving. It will go back to white if you become still again. Sometimes even if you are doing your best you will still see a red cross. This may mean that the computer is too strict and not that you are necessarily doing anything wrong. Just keep trying your best to keep the cross in the white.

The same researchers participated in data collection for all sessions. During the task, participants viewed a colored cross in the middle of the screen. FD thresholds were set to show a white cross at < 0.2 mm, a yellow cross at 0.2 mm to < 0.3 mm, and a red cross at ≥ 0.3 mm. At the conclusion of each run, participants viewed a Head Motion Report (**Supplemental Figure 1**) which displayed their performance on a gauge of 0–100 (a percentage score of 0% to 100%), and a graph of their motion level over time to help visualize total movement during the session. Participants were encouraged to bring their score closer to 100% on subsequent sequences.

### MRI acquisition and analysis

MRI data were acquired using a Siemens Prisma scanner (Siemens Medical Systems) at 3 T equipped with a 32-channel head coil. Scan sequences began with a T1-weighted structural volume using an MPRAGE sequence [repetition time (TR) = 2.4 s, echo time (TE) = 2.2 ms, flip angle = 8°, 300 × 320 matrix, voxel size = 0.8 mm isotropic]. Blood oxygenation level-dependent (BOLD) functional MRI images were acquired using a multiband echo planar imaging sequence (Feinberg et al., 2010) [TR = 3.07 s, TA = 0.770 s, TE = 37 ms, flip angle = 90º, voxel size = 2 mm isotropic, multiband factor = 8]. We used a sparse imaging design in which there was a 2.3 second delay between scanning acquisitions and the TR was longer than the acquisition time to allow for minimal scanning noise during stimulus presentation and audio recording of participant responses (Edmister et al., 1999; Hall et al., 1999). Due to slight variations in scanning sessions, participants had 763–846 frames of data (median = 794).

Analysis of the MRI data was performed using Automatic Analysis version 5.8.1 (Cusack et al., 2015) (RRID:SCR_003560) which scripted analyses run in SPM12 (Wellcome Trust Centre for Human Neuroimaging) version 7487 (RRID:SCR_007037). Functional images underwent rigid body realignment (i.e., “motion correction”). We used framewise displacement (FD) to parsimoniously summarize frame-by-frame motion, calculated by combining the six head motion estimates obtained from realignment, with a dimensional conversion of the three rotations assuming the head is a 50 mm sphere (Power et al., 2012), to produce a summary distance metric. We supplemented FD values with differential variance (DVARS), a measure of variations in image intensity. DVARS was calculated as the root-mean-squared of the time difference in the BOLD signal calculated across the entire brain, before realignment (Smyser et al., 2011). Although FD and DVARS are highly correlated, they are not identical (Jones et al., 2022), and we anticipated that these two metrics might differ in sensitivity to motion feedback.

## Results

Movement parameters for young and older adults as a function of feedback are shown in **Figure 2. Figure 2a** shows FD values for two subjects (our first young adult without feedback, and our first young adult with feedback), as well as the mean FD for all subjects in each feedback group. Although we used all time points in the statistical analysis, for illustrative purposes we also plotted summary values in **Figure 2b** (each point denoting mean FD per subject).

**Figure 2.**
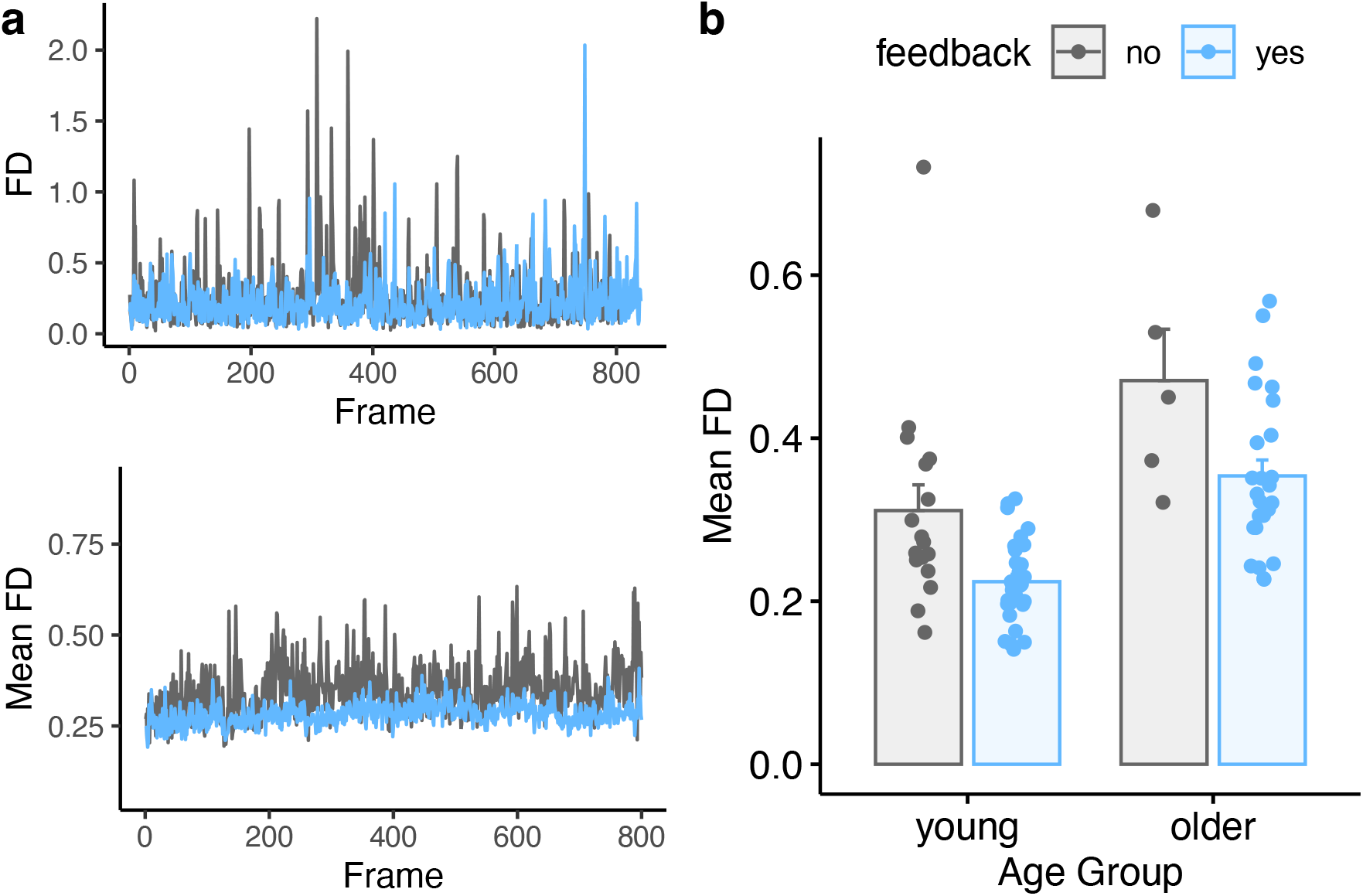
Comparison of average framewise displacement (FD) values for feedback and no-feedback groups. **a)** *Top:* FD values for two example subjects, one with feedback and one without. *Bottom:* Mean FD values for all participants in the feedback or no-feedback group. **b)** Summary of mean FD values as a function of age and feedback. Individual subjects are dots, mean ± SE shown in bars.

We analyzed these data using R version 4.2 (R Core Team, 2020) (RRID:SCR_001905). We conducted linear mixed effects analysis using the *nlme* package (Pinheiro and Bates, 2000). Because of the skewed distribution of FD data, we Gaussianized the FD values (Georg, 2011) prior to modeling using the *LambertW* package (Georg, 2022) (see **Supplemental Figure 1**).

We accounted for temporal autocorrelation (that is, motion at one frame is related to motion during the following frame) using a first-order autoregressive model. The model specification was:

~~~
m2 <-lme(FD ∼ 1 + frame + age_group * feedback,
  random = ∼ frame | subject_number,
  data = df,
  method = “ML”,
  correlation = corAR1(value =
autocorrelation_estimate, form = ∼ frame | subject_number, fixed
= FALSE))
~~~

where `df` is the data frame, `FD` is framewise displacement, and `feedback` is a dichotomous variable denoting whether feedback was provided or not (reference: no feedback). We calculated the `autocorrelation_estimate` from the data (value of 0.55). Model results are shown in **Table 2**. Of note is a significant effect of feedback.

**Table 2.** Fixed effects results for motion (FD) model by frame.

|  | Est | SE | df | t | p |  |
| --- | --- | --- | --- | --- | --- | --- |
| Intercept | 0.3572638 | 0.021880945 | 61778 | 16.32762<br>1 | < .0001 | *** |
| frame | 0.0000273 | 0.000008824 | 61778 | 3.088461 | .0020 | ** |
| age group | -0.1033003 | 0.024837814 | 74 | -4.158994 | 0.0001 | *** |
| feedback | -0.0534164 | 0.023917879 | 74 | -2.233326 | 0.0286 | * |
| age group x feedback | 0.0184424 | 0.028091970 | 74 | 0.656502 | 0.5135 |  |

\*\*\* $p < .001$ ; \*\* $p < .01$ ; \* $p < .05$

Although we first included time (frame) in the model, in fMRI research it is often convenient to think about continuous scanning runs (sometimes also referred to as “sessions”). In the current study, each participant had 6 runs of data. We thus ran an additional model to see whether the effect of feedback was comparable across run. Because time was expressed as 6 runs, we did not account for temporal autocorrelation in this model:

~~~
m3 <-lme(FD ∼ 1 + run + age_group * feedback,
   random = ∼ run | subject_number,
   data = df,
   method = “ML”,
   correlation = NULL)
~~~

Results of this analysis by run are shown in **Table 3**, and FD as a function of run are shown in **Figure 3**. We again found a significant effect of feedback.

**Table 3.** Fixed effects results for motion (FD) model by run.

|  | Est | SE | df | t | p |  |
| --- | --- | --- | --- | --- | --- | --- |
| Intercept | 0.3555292 | 0.020471649 | 61774 | 17.366906 | < .0001 | *** |
| run2 | 0.0111638 | 0.004760753 | 61774 | 2.344960 | 0.0190 | * |
| run3 | 0.0189044 | 0.005247816 | 61774 | 3.602345 | 0.0003 | ** |
| run4 | 0.0229300 | 0.005437600 | 61774 | 4.216932 | < .0001 | *** |
| run5 | 0.0206506 | 0.006320480 | 61774 | 3.267252 | 0.0011 | ** |
| run6 | 0.0179240 | 0.006190537 | 61774 | 2.895389 | 0.0038 | * |
| age group | -0.1040961 | 0.023205030 | 74 | -4.485930 | < .0001 | *** |
| feedback | -0.0605921 | 0.022347092 | 74 | -2.711407 | 0.0083 | ** |
| age group x feedback | 0.0261700 | 0.026245629 | 74 | 0.997117 | 0.3220 |  |

\*\*\* p < .001; \*\* p < .01; \* p < .05

**Figure 3.**
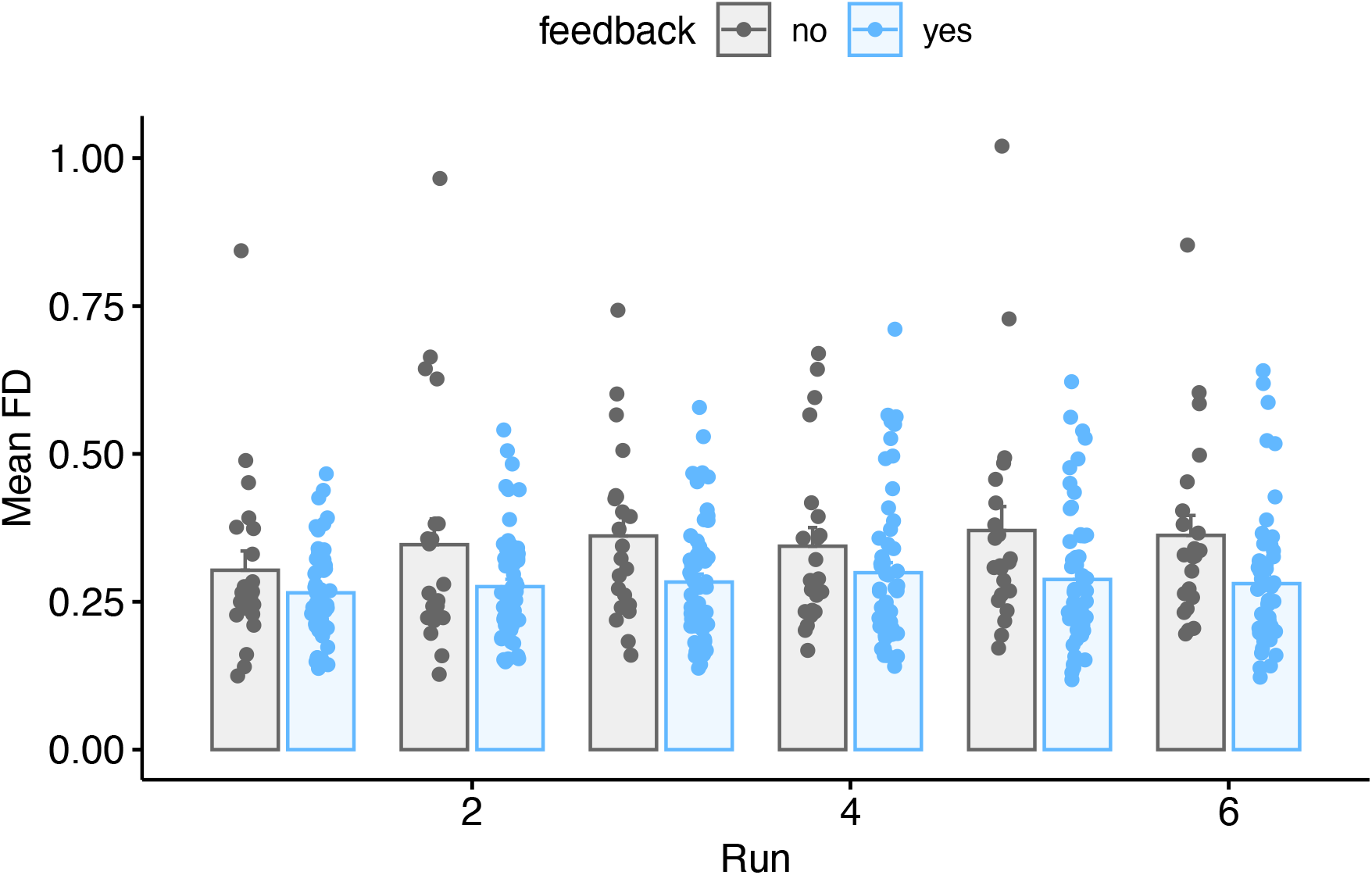
Mean FD values for all participants in the feedback or no-feedback group as a function of run. Individual subjects are dots, mean ± SE shown in bars.

Even though our analysis was focused on changes in mean FD, we wanted to look in more detail at how feedback affected the distribution of movement. We thus plotted the density of FD values for all scans as a function of feedback (**Figure 4**) for FD values up to 2.0. We used density rather than the count to control for the different numbers of frames in the two groups (resulting from different numbers of subjects). Although qualitative, this analysis highlights that participants receiving feedback showed fewer high-FD scans than those receiving lower feedback, with this effect becoming more apparent at higher FD values—for FD values above 1.5, the feedback group had less than a third of these scans than the group without feedback.

**Figure 4.**
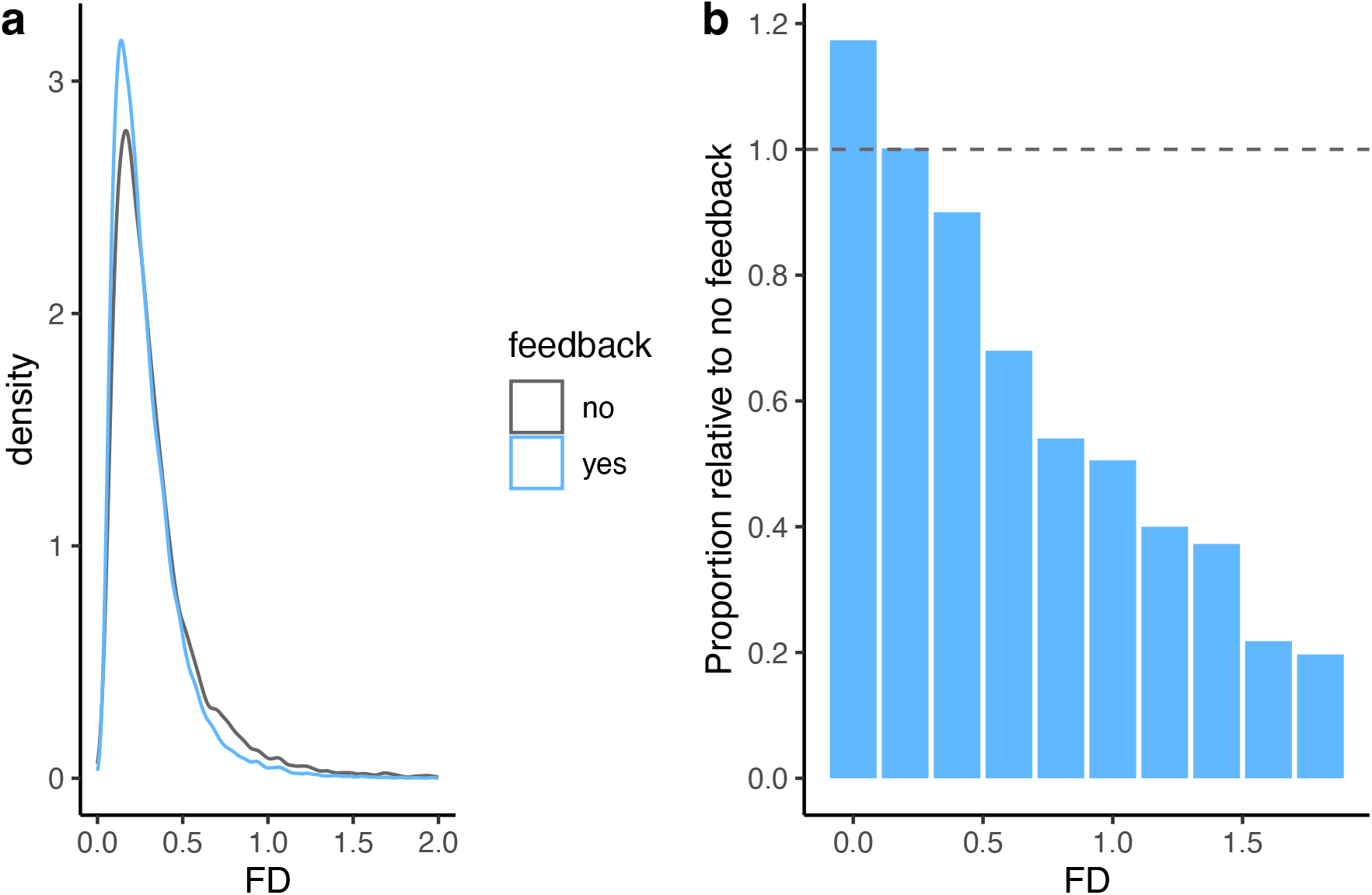
**a)** Density plot of all FD values as a function of feedback. **b)** Density of FD values in the feedback condition relative to the no feedback condition. A value of 1 (dashed line) indicates equal density of FD values in the two conditions. The plot illustrates a decrease in high-FD values in the feedback condition, and an increase in low FD values, relative to the no feedback condition.

We next repeated our primary FD analyses, but using DVARS (rather than FD) as a dependent measure, under the assumption that FD and DVARS may be differently sensitive to effects of motion, and one or the other may be of more interest in any specific study. As with FD, we used Gaussianized data to reduce the influence of the skewed distribution. Summary results are shown in **Figure 5** and **Table 4**. Results as a function of run are additionally shown in **Figure 6** and **Table 5**. Although DVARS was numerically lower for the feedback group than the no-feedback group, the difference was not statistically significant.

**Table 4.** Fixed effects results for DVARS model by frame.

|  | Est | SE | df | t | p |  |
| --- | --- | --- | --- | --- | --- | --- |
| Intercept | 228.48139 | 9.306936 | 61778 | 24.54958<br>4 | <.0001 | *** |
| frame | 0.00521 | 0.003254 | 61778 | 1.602724 | 0.1090 |  |
| age group | -24.05534 | 10.517516 | 74 | -2.287169 | 0.0250 | * |
| feedback | -15.40664 | 10.127865 | 74 | -1.521213 | 0.1325 |  |
| age group x feedback | 2.17760 | 11.895410 | 74 | 0.183062 | 0.8552 |  |

\*\*\* $p < .001$ ; \*\* $p < .01$ ; \* $p < .05$

**Table 5.** Fixed effects results for DVARS model by run.

|  | Est | SE | df | t | p |  |
| --- | --- | --- | --- | --- | --- | --- |
| Intercept | 227.31594 | 8.962557 | 61774 | 25.36284<br>3 | < .0001 | *** |
| run 2 | 2.05963 | 1.436412 | 61774 | 1.433874 | 0.1516 |  |
| run 3 | 3.98708 | 1.706406 | 61774 | 2.336538 | 0.0195 | * |
| run 4 | 5.40560 | 1.858504 | 61774 | 2.908578 | 0.0036 | ** |
| run 5 | 4.34756 | 2.478246 | 61774 | 1.754289 | 0.0794 |  |
| run 6 | 2.73375 | 2.081469 | 61774 | 1.313377 | 0.1891 |  |
| age group | -23.11906 | 10.120695 | 74 | -2.284335 | 0.0252 | * |
| feedback | -16.27255 | 9.745900 | 74 | -1.669682 | 0.0992 |  |
| age group x feedback | 2.64483 | 11.446654 | 74 | 0.231057 | 0.8179 |  |

\*\*\* $p < .001$ ; \*\* $p < .01$ ; \* $p < .05$

**Figure 5.**
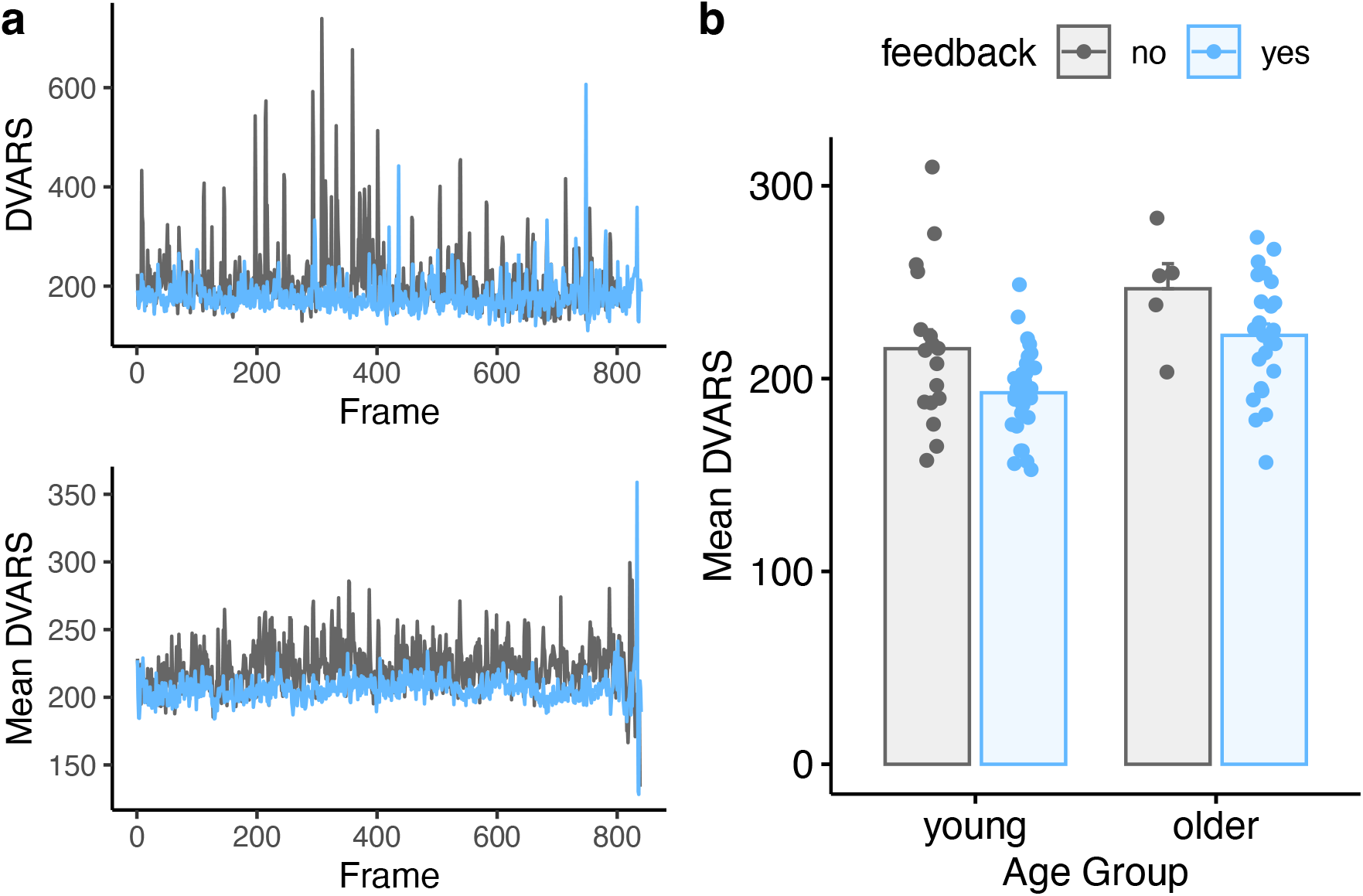
Comparison of average DVARS values for feedback and no-feedback groups. **a)** *Top:* DVARS values for two example subjects, one with feedback and one without. *Bottom:* Mean DVARS values for all participants in the feedback or no-feedback group. **b)** Summary of mean DVARS values as a function of age and feedback. Individual subjects are dots, mean ± SE shown in bars.

**Figure 6.**
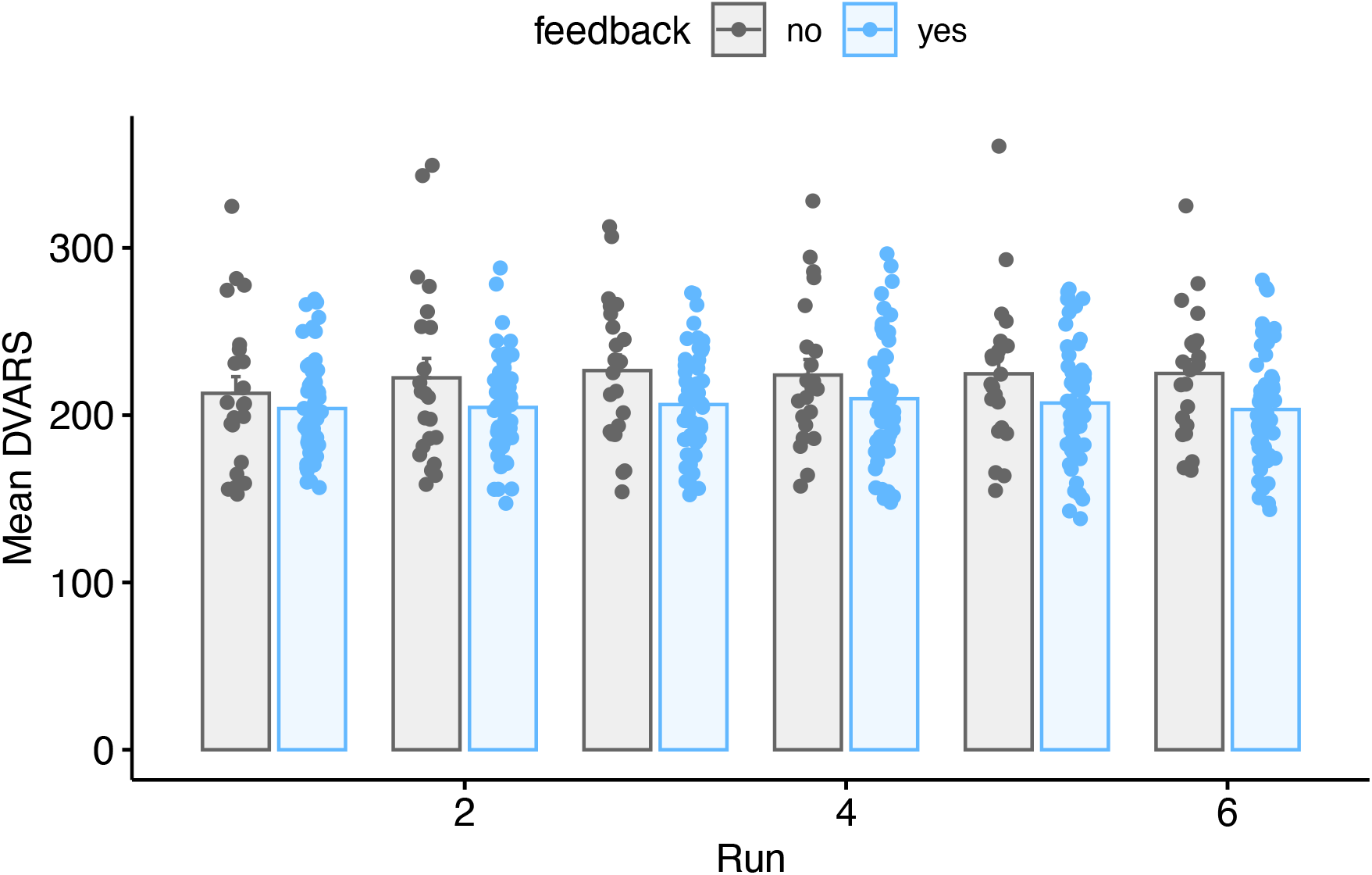
Mean DVARS values for all participants in the feedback or no-feedback group as a function of run. Individual subjects are dots, mean ± SE shown in bars.

In addition to framewise measurements of FD and DVARS, we investigated whether motion feedback affected the temporal signal-to-noise ratio (tSNR) of the data. We first created a brain mask using each subject’s tissue class segmentation, binarizing at a gray matter probability of 0.8. We then calculated tSNR (the mean of the signal divided by its standard deviation, over frames) in every voxel within the brain mask. We then performed two complementary analyses. First, for each run, we took the mean tSNR over voxels, and used the same statistical model as for FD and DVARS, shown in **Figure 7** and shown in **Table 6**. Although the tSNR values were numerically higher in the feedback group, there was no significant effect of feedback (p = 0.0814).

**Table 6.** Fixed effects results for tSNR model.

|  | Est | SE | df | t | p |  |
| --- | --- | --- | --- | --- | --- | --- |
| Intercept | 35.62189 | 2.3494343 | 389 | 15.161899 | < .0001 | *** |
| run | -0.01283 | 0.1116692 | 389 | -0.114888 | 0.9086 |  |
| age group | 5.61719 | 2.6542796 | 74 | 2.116276 | 0.0377 | * |
| feedback | 4.51521 | 2.5559386 | 74 | 1.766556 | 0.0814 |  |
| age group x feedback | -0.11883 | 3.0020112 | 74 | -0.039584 | 0.9685 |  |

\*\*\* $p < .001$ ; \*\* $p < .01$ ; \* $p < .05$

**Figure 7.**
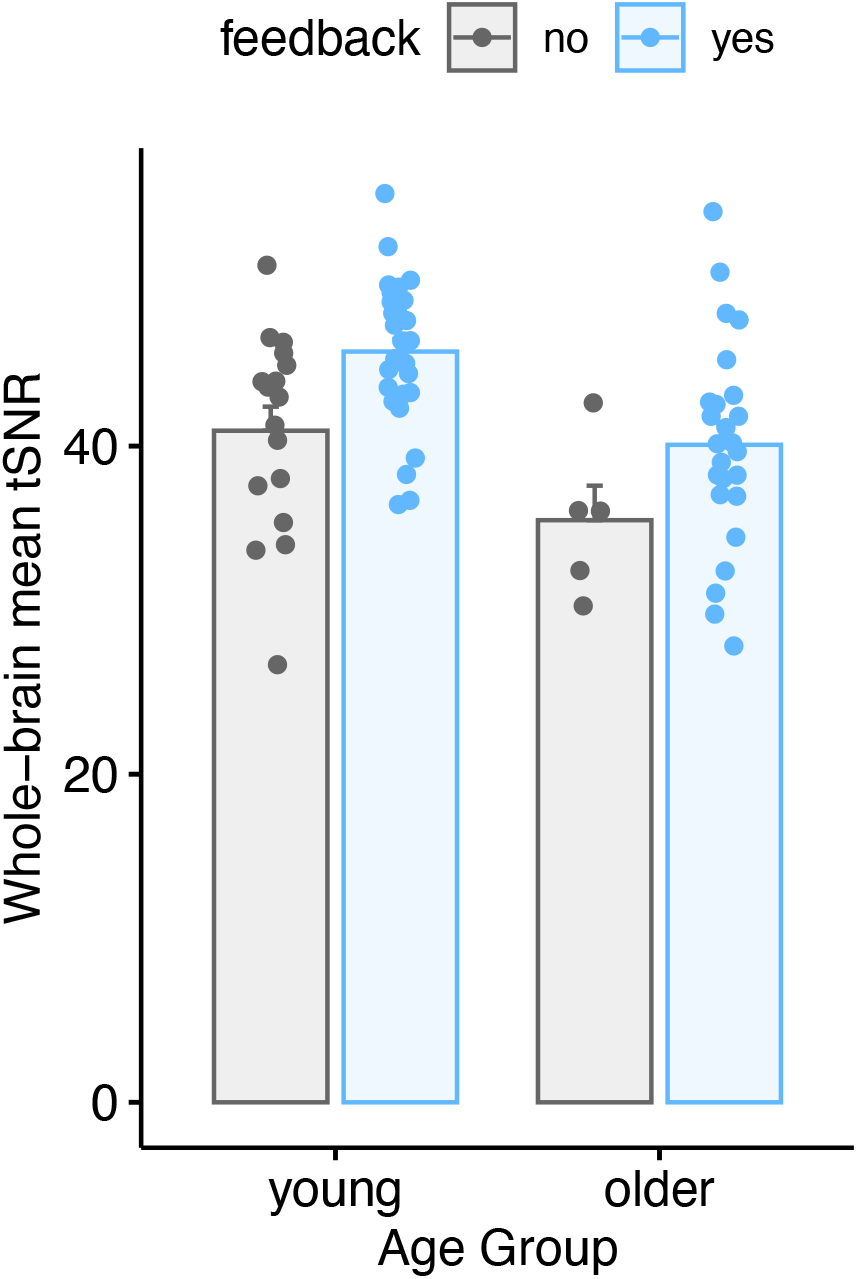
Comparison of average temporal SNR (tSNR) values for feedback and no-feedback groups. Each participant is represented by a single point per run, averaging the tSNR over all voxels in the gray matter mask.

Finally, we assessed the degree to which feedback may have affected performance on the behavioral task. Accuracy on the word repetition task is shown in **Figure 8** as a function of age group and feedback. We used a linear model to test for the impact of feedback on behavioral accuracy (averaging over experimental runs). We found no significant effect of age group, no significant effect of feedback, and no significant age x feedback interaction, although in older adults there was a small numerical decrease in accuracy during the feedback condition.

**Figure 8.**
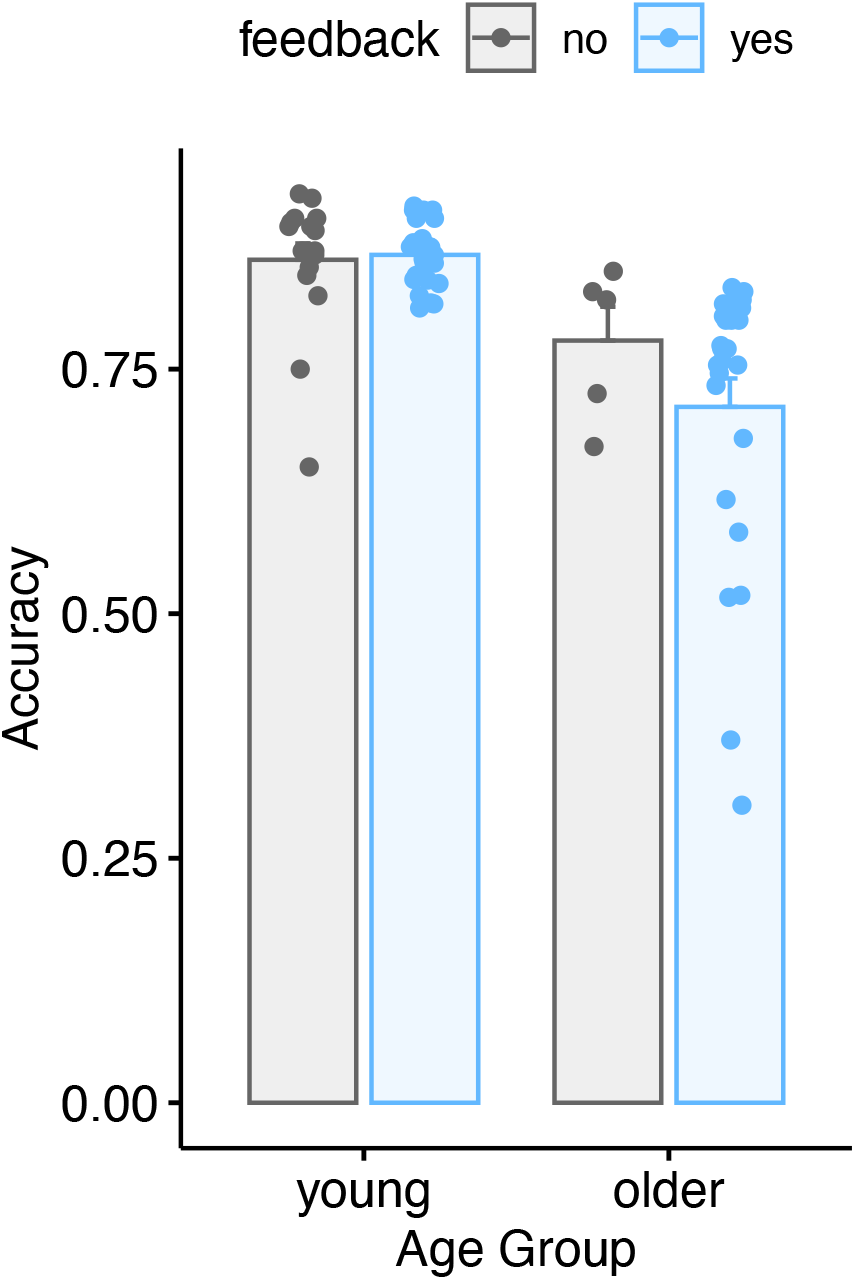
Behavioral accuracy for young and older adults as a function of feedback.

## Discussion

Given the pernicious and sometimes unpredictable effects of head movement in fMRI studies, reducing motion in the first place is on its face an appealing strategy. In the current study we evaluated the degree to which a combination of real-time and summary motion feedback could reduce head movement in adult participants by pseudorandomly assigning participants to feedback or no feedback groups. We found that motion feedback was associated with a statistically significant reduction in average FD of approximately 0.06, with consistent (though not significant) changes in DVARS and tSNR. We discuss these findings and their implications below.

First, our primary finding that providing motion feedback can significantly reduce head motion in participants undergoing task-based MRI is notable. Our findings are also consistent with those of Yang and colleagues (2005), who reported a reduction in head motion after integrating visual feedback in the display of an N-back task. These findings add motion feedback to the limited set of tools MRI researchers have for reducing head motion during acquisition. Although numerous strategies exist for accounting for head motion in preprocessing and analysis, none are perfect, and it is difficult to ascertain the most appropriate approach (Ardekani et al., 2001; Oakes et al., 2005; Johnstone et al., 2006; Jones et al., 2022). Reducing motion during acquisition may therefore be a preferable alternative in many circumstances.

It is important to highlight the magnitude of the effect we observed. Compared to the participants with no motion feedback, participants with feedback showed a reduction in FD of approximately .06. Although this may be helpful in reducing motion-related artifacts in data processing, it is not a large effect (though in theory it might be increased if different types of feedback were used, or in the context of a different task). It is also important to remember that we tested healthy adults aged 19–81. It could be that the effect of motion feedback would be larger (or smaller) in different populations.

That being said, the principal effect of feedback may not be the *average* reduction in FD observed, but rather a reduction in the number of large motion events. Rapid head motion tends to have a larger effect on the BOLD signal than slower motion. Thus, reducing the number of large, quick movements may improve signal quality to a greater extent than a simple reduction in mean FD. Indeed, we saw the biggest proportional difference between feedback groups at higher FD values (**Figure 4b**). Interestingly, the reduction in high motion events roughly corresponds to an FD of 0.3, the value at which participants received “red” feedback.

An important consideration when implementing real-time feedback during fMRI is whether doing so changes task demands. Although we found accuracy was statistically comparable across feedback groups, we cannot rule out other changes. Indeed, we observed a numeric (albeit not significant) decrease in task accuracy in older adults when they were presented with real-time feedback relative to the no-feedback group. Broadly speaking, we might expect monitoring the display for feedback and adjusting one’s movement in response to engage systems related to cognitive control (sometimes also referred to as attentional control or executive function) (see also McCabe et al., 2010). In one framing of cognitive control, cognitive control is proposed to operate in at least two complementary modes: a *proactive* mode that is concerned with maintaining goal-directed behavior, and a *reactive* mode that is engaged when goal-directed performance falters (Braver, 2012). There are two implications of such a framework. First, participants who are less able to engage cognitive control may benefit less from motion feedback compared to those who are better able to engage control systems. Second, researcher concerns about how task demands may affect imaging results of interest may depend on the nature of the main fMRI task. For example, tasks tapping cognitive control or executive function may be more impacted by motion feedback than those focused on primary sensory or motor systems. In our case, we used a speech-in-noise task, one that is hypothesized to engage cognitive control networks (Peelle, 2018; McLaughlin et al., 2021).

We did notice a trend towards age effects, such that older adults’ behavioral performance was more affected than that of young adults in the feedback condition. In this context it may also be relevant that older adults can be more sensitive to errors (and thus differentially sensitive to feedback). For example, in drift diffusion models of cognitive tasks, older adults often differ in threshold than drift rate compared to young adults (e.g., Ratcliff et al., 2004).

In our implementation, participants received brief instructions before entering the scanner, and once in the scanner the display was explained to them. One interesting future direction would be to provide participants with motion training over a longer period to see whether they might learn to better limit their motion in the absence of feedback (for example, by training in a mock scanner). Such an approach would take more time, but might circumvent some of the challenges associated with real-time feedback (such as introducing a “dual-task” situation) (Pashler, 1994).

Although we found a significant effect of motion feedback on FD, we did not observe a significant effect on motion feedback on DVARS (although DVARS values were numerically lower in the feedback group). FD and DVARS are typically strongly correlated (Jones et al., 2022), but not identical, and thus the divergence in findings is not entirely unexpected. The numerical (albeit not significant) reductions in DVARS in participants receiving motion feedback is consistent with the effect we saw for FD. In addition, we found that feedback led to increased whole-brain tSNR, which we used as a proxy of overall image quality. Despite the tSNR being numerically larger in the group receiving motion feedback, the difference was not statistically significant. However, it is still possible that small improvements in tSNR would result in appreciable improvements in the accuracy or reliability of statistical models.

There are several important caveats associated with the current report. One is that the instructions were not perfectly matched across group; specifically, the group receiving feedback was told that “[e]ven very small movements that you are not aware of can affect the image quality”, an instruction not given to the group not receiving feedback. It could be that the specific wording of the instructions affected participant behavior. A related point pertains to our task design. We opted for a between-subjects design to avoid any “contamination” of motion feedback. However, this choice necessarily resulted in groups that contained different individuals (and in our case, different instructions). An alternative design would be a within-subjects manipulation, which may be a fruitful area for further extensions of this work.

Finally, it is worth considering real-time feedback using estimated motion, as implemented here, compares with other real-time feedback approaches. One particularly attractive approach was introduced by Krause and colleagues (2019), in which the authors affixed medical tape to participants’ heads. The adhesion of the tape provided tactile feedback when participants moved their heads, and this real-time feedback significantly reduced participant’s head motion (both translation and rotation). An added advantage of this approach is the simplicity of implementation and lack of custom software (e.g., real-time analysis of movement parameters) or interference with task (i.e., tactile feedback does not interfere with visual stimuli). Individual researchers will need to decide which approach to motion reduction, if any, is appropriate for a given task and specific population.

In conclusion, we found that motion feedback as implemented in FIRMM significantly reduced the amount of head motion we observed during task-based fMRI. Real-time feedback may thus be well-suited to complement other approaches to motion reduction depending on the specific needs of a given study.

## Data availability statement

The data that support the findings of this study are openly available in OpenNeuro at https://doi.org/10.18112/openneuro.ds004285.v1.0.0, reference number ds004285.

## Ethics statement

The study reported here was conducted according to a protocol approved by the Washington University in Saint Louis Institutional Review Board.

## Acknowledgments

Work reported here was supported by grant R01 DC014281 from the US National Institutes of Health. Support was also provided by the Basque Government through the BERC 2022-2025 program and by the Spanish State Research Agency through BCBL Severo Ochoa excellence accreditation CEX2020-001010-S. The multiband echo planar imaging sequence was provided by the University of Minnesota Center for Magnetic Resonance Research. We are grateful to Linda Hood for assistance with data collection, and to Nico Dosenbach and Jackie Hampton for assistance implementing FIRMM.

## Footnotes

1 Now a commercial product offered by Turing Medical (St. Louis, MO).

