## Supplemental for "Real-time feedback reduces participant motion during task-based fMRI"

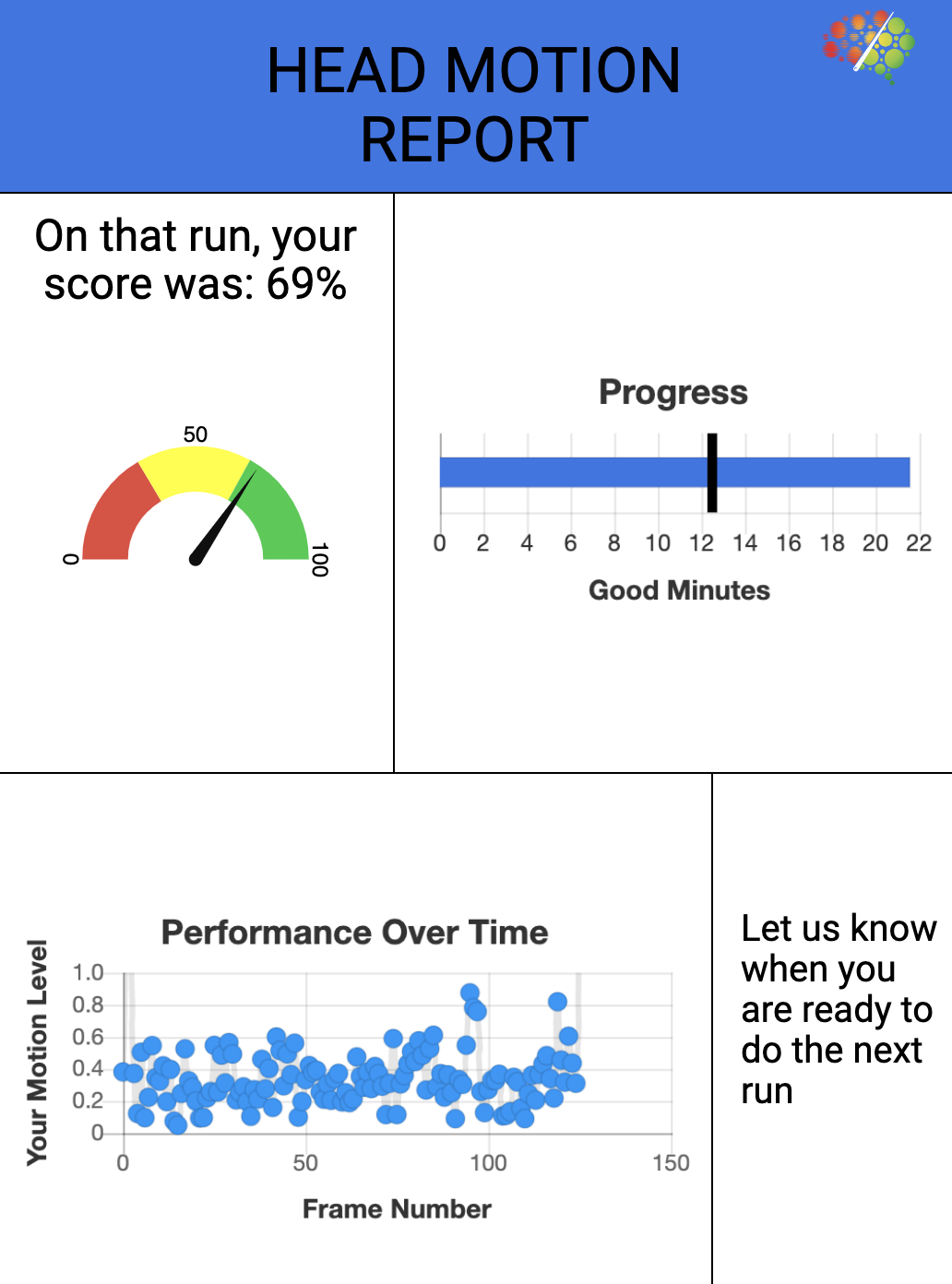

**Supplemental Figure 1.** Head motion report provided to participants following each run of the experiment. The upper right quadrant indicates how many minutes of data with acceptable motion has been collected (under the logic that in a typical resting state paradigm, additional data might be collected to meet a threshold for amount of good data) (Dosenbach et al., 2017). The percentage score relates to how much of the time the participant had motion in an acceptable range.

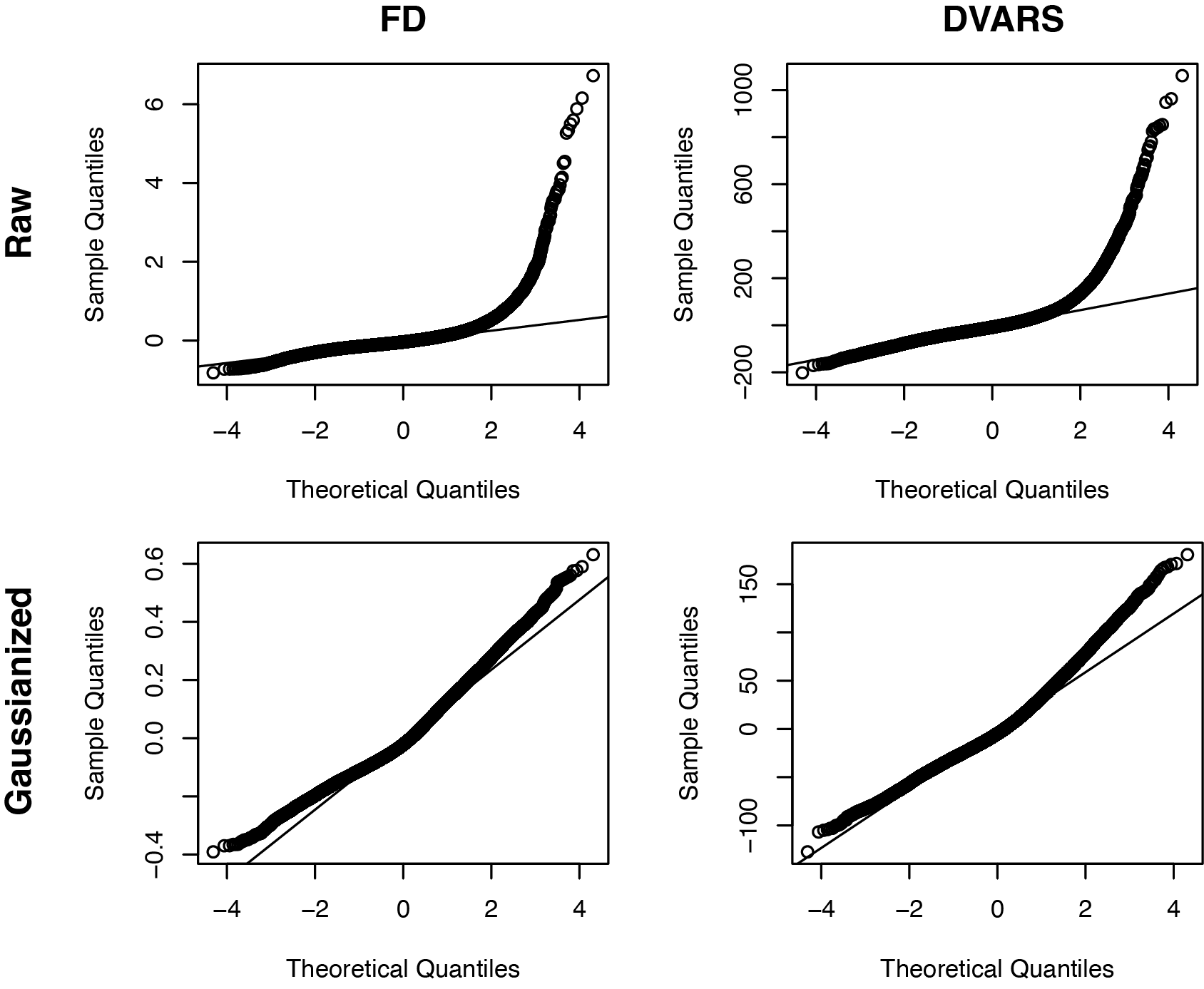

**Supplemental Figure 2.** Q-Q plots of model residuals for FD and DVARS models with raw data (top row) and Gaussianized data (bottom row).
